# CAMUS: Scalable Phylogenetic Network Estimation

**DOI:** 10.64898/2026.02.01.703143

**Authors:** James Willson, Tandy Warnow

## Abstract

**Motivation:** Phylogenetic networks are models of evolution that go beyond trees, and so represent reticulate events such as horizontal gene transfer or hybridization, which are frequently found in many taxa. Yet, the estimation of phylogenetic networks is extremely computationally challenging, and nearly all methods are limited to very small datasets with perhaps 10 to 15 species (some limited to even smaller numbers).

**Results:** We introduce CAMUS (Constrained Algorithm Maximizing qUartetS), a scalable method for phylogenetic network estimation. CAMUS takes an input constraint tree *T* as well as a set *Q* of unrooted quartet trees that it derives from input, and returns a level-1 phylogenetic network *N* that is built upon *T* through the addition of edges, in order to maximize the number of quartet trees in *Q* that are induced in *N* . We perform a simulation study under the Network Multi-Species Coalescent and show that a simple pipeline using CAMUS provides high accuracy and outstanding speed and scalability, in comparison to two leading methods, PhyloNet-MPL used with a fixed tree and SNaQ. CAMUS is slightly less accurate than PhyloNet-MPL used without a fixed tree, but is much faster (minutes instead of hours) and can complete on inputs with 201 species while PhyloNet-MPL fails to complete on the inputs with more than 51 species.

**Availability and Implementation:** The source code is available at https://github.com/jsdoublel/camus.

## 1 Introduction

Phylogenetic trees form an important basis for biological discovery, including understanding adaptation and biodiversity. The estimation of phylogenetic trees for a single gene can be fairly simple, but estimations of phylogenetic trees for multiple genes (i.e., species trees), and especially for genome-scale data, is complicated by processes such as incomplete lineage sorting (ILS) and gene duplication and loss (GDL), in which different genomic regions (“genes”) have different trees [Maddison, 1997]. Much progress has been made on estimating species trees and new methods capable of high accuracy are now available (e.g., Mirarab et al. [2014], Zhang et al. [2020]) and in wide use in biological studies.

However, when processes such as hybrid speciation or horizontal gene transfer occur, the tree model is no longer appropriate, and instead a phylogenetic network, which can be seen as a tree with extra edges (and so has cycles when considered as undirected graphs), is needed [Morrison, 2014, Huson and Bryant, 2006, Kong et al., 2025].

Several methods have been developed for estimating phylogenetic networks, which differ by approach, the evolutionary processes they consider, and the constraints they make on the network topology. For example, Gusfield [2005] examined a population-genetics context where each site evolves down a rooted tree contained within the rooted phylogenetic network under the infinite sites assumption with HGT but not ILS [Gusfield, 2005, Warnow et al., 2025]. Other researchers have addressed phylogenetic network estimation under the Network Multi-Species Coalescent (NMSC) (see discussion in Mirarab et al. [2021]), which models both HGT and ILS.

Methods for estimating phylogenetic networks are challenged by statistical and computational issues. Specifically, when phylogenetic networks can have cycles that share vertices, Gambette and Huber [2012] showed that two different networks can have the same rooted triplet trees. In contrast, rooted phylogenetic networks in which no two cycles share nodes and all cycles are large enough are uniquely defined by their rooted triplets [Gambette and Huber, 2012]. For this reason, a substantial focus has been made on developing methods for estimating phylogenetic networks where the cycles are vertex-disjoint, which are referred to as level-1 phylogenetic networks or “galled trees” (see Gusfield [2005] for an early paper on level-1 phylogenetic network estimation).

Methods for estimating level-1 phylogenetic networks from gene trees under the NMSC have been developed, several of which are provably statistically consistent provided that there are no small cycles (e.g., various tools within the PhyloNet package [Than et al., 2008], SNaQ [Solís-Lemus and Ané, 2016], and NANUQ+ [Allman et al., 2019]). Of these, PhyloNet-MPL [Yu and Nakhleh, 2015] and SNaQ, both of which optimize maximum pseudolikelihood, are probably the most frequently used tools for level-1 phylogenetic network analysis, and have been shown to be able to run on datasets with at least 10 species. However, even these methods typically have very large computational requirements, often using many hours to complete on inputs with only 10-20 species, and failing to complete on inputs with just 50 species using available resources.

An alternative approach to phylogenetic network estimation operates in two-phases: first a rooted tree is computed, and then edges are added to the tree to create a phylogenetic network. While such approaches may fail to provide a guarantee of statistical consistency (especially in the case of anomalous networks, see Ané et al. [2024]), they can be fast and highly scalable. The approach we present here falls into this category of technique.

We present CAMUS (Constrained Algorithm Maximizing qUartetS) as well as pipelines using CAMUS designed to be used for level-1 phylogenetic network estimation from sequence data or gene trees. For example, when estimating a phylogenetic network under the NMSC, the input is a set of estimated gene trees. Given this input, in the first step we compute a rooted species tree *T* and also a set Q of quartet trees (i.e., unrooted trees on four species). Then, CAMUS adds edges to *T*, producing a rooted level-1 phylogenetic network that maximizes the number of quartet trees in Q induced by rooted trees within the network. CAMUS uses a dynamic programming approach that solves the optimization problem exactly in polynomial time. We also present a technique for selecting the quartet trees for Q from the estimated gene trees, and show that using this set for Q provides high accuracy when using CAMUS to produce a level-1 phylogenetic network.

Our study on datasets with up to 201 species and 1000 genes establishes that CAMUS has substantial advantages over both SNaQ and PhyloNet-MPL, each of which was given the same tree to use either as a constraint tree (for CAMUS and one way of running PhyloNet-MPL) or starting tree (for SNaQ and the other way of running PhyloNet-MPL). CAMUS was far more scalable than other methods. When limited to 12 hours and 256 GB of memory, CAMUS was the only method that was able to complete on the 201-species dataset, while the next most scalable method—PhyloNet-MPL used with the same starting tree as a constraint—timed out on the datasets with more than 51 species. The other methods, SNaQ and PhyloNet-MPL without a fixed starting tree, were even less scalable.

CAMUS was also more accurate than SNaQ and PhyloNet-MPL used with a constraint tree. PhyloNet-MPL run without a constraint tree achieves slightly better accuracy than CAMUS, but incurs a substantial computational hit, completing on only the smaller datasets in the allotted time (taking more than 3.5 hours on the 51-species datasets to add just one reticulation, where CAMUS finishes in under a minute and adds multiple extra reticulations).

## 2 CAMUS

### 2.1 Preliminary material

We begin with some terminology and basic observations.

Each rooted binary level-1 phylogenetic network *N* induces a set *𝒯* of rooted binary trees formed by deleting one incoming edge into each reticulation node (i.e., nodes that have indegree 2) in each of the possible ways. Given tree *t* ∈ *𝒯*, we consider *t* in its unrooted form and define the set *Q*(*t*) of (unrooted) quartet trees induced in *t*. We define *Q*(*N*_*r*_) = U_*t*∈*𝒯*_ *Q*(*t*); the use of the subscript *r* indicates its reliance on the rooting of *N*, and distinguishes it from *Q*(*N*), which is the set of all unrooted quartet trees found in spanning trees of the unrooted version of *N* (see discussion in Warnow et al. [2025]).

#### Definition 1.

*Given* 𝒬, *T, and N, where N is a level-1 phylogenetic network built upon T, we say that q* ∈ 𝒬 *is* ***satisfied by* N** *if q* ∈ *Q*(*N*_*r*_) \ *Q*(*T*). *We define* **T** + **e**, *where e is a non-tree edge in N, to be the subnetwork of N defined by removing all other non-tree edges and then suppressing nodes that have indegree and outdegree one. Furthermore, if N* = *T* + *e then we also say* **q *is satisfied by* e**. *These concepts are illustrated in Figure 1*.

**Figure 1:**
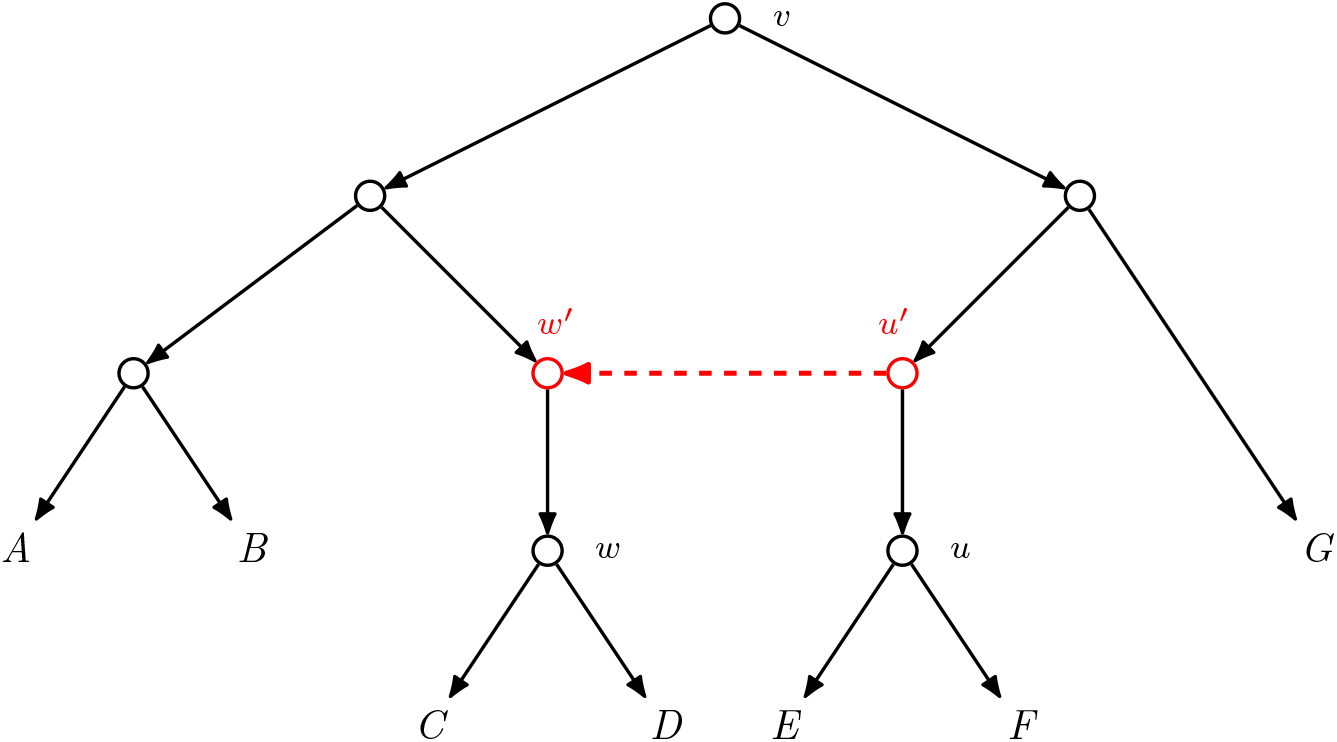
Tree-based network *N* with underlying tree *T* defined by solid black edges and one non-tree directed edge (dashed red). *N* can be constructed by adding the non-tree edge to *T*, and we say that the non-tree edge is “above the ordered pair (*u, w*)”, which means that the edges above *u* and *w* are subdivided, creating two new nodes (here denoted by *u*^*′*^ and *w*^*′*^), and a directed edge is then inserted from *u*^*′*^ to *w*^*′*^. *N* has a single cycle, which we denote by *γ*_(*u,w*)_. The unique node with indegree two is *w*^*′*^, which we refer to as the bottom node of the cycle. We also refer to this network as *T* + *e*, where *e* is the non-tree edge. Note that *N* induces two rooted trees on *A, D, E, G*: the rooted tree ((*A, D*), (*E, G*)) (which is also induced by *T*) and the rooted trees (*A*, ((*D, E*), *G*)) (which does not appear in *T*). The rooted four-leaf trees define two unrooted trees on the leafset, referred to as quartet trees: *AD | EG* and *DE | AG*. We say that the quartet tree *DE | AG* is satisfied by *N* but not by *T*, and we also say it is satisfied by the added edge from *u*^*′*^ to *v*^*′*^.

#### Lemma 1.

*For any rooted binary tree T and extension of T into level-1 phylogenetic network N, if quartet tree q is satisfied by N, then there is a unique non-tree edge e such that q is satisfied by T* + *e*.

*Thus, the set of quartets that are satisfied by N is the disjoint union of quartets satisfied by T* + *e, as we let e vary among the non-tree edges of N*.

*Proof*. Proof by contradiction. Suppose there are two edges in *N* that satisfy a quartet *q*. Since *q* ∉ *Q*(*T*_*r*_), there are two vertex-disjoint cycles used to induce *q*, and the four leaves of *q* attach to four different nodes in each cycle (see Warnow et al. [2025]). Let *v* be the LCA of the roots of the two cycles. Then *v* is in a cycle that contains the roots of both cycles, and *N* is not level-1. ◻

Note that *T* + *e* contains the endpoints of edge *e*, but these endpoints are not in *T* ; instead, these are *added* to *T* by subdividing the relevant edges (see Figure 1). We use this observation in the following:

#### Definition 2.

Procedure ASSIGN *Given N, T, and quartet q satisfied by N and not by T, find* (*u*^′^, *w*^′^), *the unique non-tree (directed) edge that satisfies q. Find u and w in T directly below u*^′^ *and w*^′^, *respectively, and let v* = *lca*_*T*_ (*u, w*). *Assign q and the non-tree edge* (*u*^′^, *w*^′^) *satisfying q to the triplet* (*u, w, v*). *Denote v as the “target” of q and its satisfying non-tree edge. Figure 1 illustrates* Procedure ASSIGN, *with the red non-tree edge and quartet DE*|*AG both assigned to the triplet* (*u, w, v*). .

We continue with some observations about this relationship between non-tree edges and target nodes in level-1 phylogenetic networks.

#### Lemma 2.

*Let N be a level-1 phylogenetic network N extending T, and let v be a node in T* . *Then there is at most one non-tree edge in N for which v is the target*.

*Proof*. If there were two non-tree edges in *N* with *v* the target, then *N* would contain two cycles, and it is easy to see that each would contain *v* as a node, and hence the two cycles would not be disjoint. This violates *N* being level-1. ◻

We continue by defining some necessary terms, and then provide the DP algorithm for CAMUS.

#### Definition 3.

*Let T be a rooted tree and u and w be nodes in N. The phrase* ***add edge above*** (**u, w**) *means we create two new vertices in T—one bisecting the edge above u and one bisecting the edge above w—then creating an edge from the vertex above u to the vertex above w (see Figure 1). This creates a network N that has a cycle, denoted by* ***γ*** (**u, w**), *for which the vertex created above w will be the bottom node of the cycle (i*.*e*., *the only node in the cycle with indegree* 2).

#### Definition 4.

*Let T be a rooted tree and let q be a quartet tree. We define* ***δ***_**T**_(**q**, (**u, w**)) *to be the number of times q appeared in* 𝒬 *if q is satisfied by an edge e added above* (*u, w*); *otherwise, the value is* 0.

#### Definition 5.

*Let N be a rooted level-1 phylogenetic network and v a node in N. Then* **l**(**v**) *refers to the left child of v and* **r**(**v**) *refers to the right child of v*.

### 2.2 The DP formulation

The high-level approach in CAMUS is that we process the constraint tree *T* from the leaves upward, filling out a dynamic programming (DP) array *M* that has entries for every node *v* in *T*, so that *M* [*v*] is equal to the largest number of quartets in 𝒬 that can be satisfied by a level-1 phylogenetic network produced by adding edges that have target nodes in *T*_*v*_ . Once this array is filled in, the solution is obtained at the root, and backtracking allow us to infer the optimal network.

Consider the following quantity:

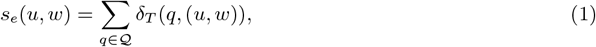

which is the number of quartets in 𝒬 satisfied by the edge *e* added above (*u, w*) (see Definition 4). Because, in a level-1 phylogenetic network *N*, each quartet that is satisfied by *N* will be satisfied by exactly one added edge (Lemma 1). it follows that *M* [*v*] will be the maximum sum of *s*_*e*_(*u, w*), as (*u, w*) ranges over a set of added edges whose target nodes are in *T*_*v*_ where the set together defines a level-1 phylogenetic network. We therefore calculate all these variable *s*_*e*_(.) in a preprocessing step, as they do not depend on other calculations.

We now present the DP algorithm. The base case is simply that *M* [*v*] = 0 for all *v* where *v* is a leaf. Next, we fill out the dynamic programming matrix from the bottom up:

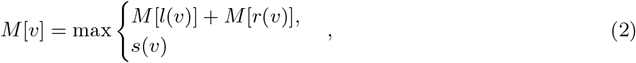

where *s*(*v*) (formally defined below) represents the case where the optimal solution is achieved by adding an edge whose target node is *v* (see Definition 2). Note that the first part of the assignment, where *M* [*v*] = *M* [*l*(*v*)] + *M* [*r*(*v*)], handles the case where the optimal score is obtained without adding any edge whose target node is *v*.

We now define *s*(*v*). We begin by defining *Pairs*(*v*), a set of ordered pair (*u, w*) of nodes in *T* so that the triplet (*u, w, v*) would be assigned to a non-tree edge we would add to *T* (see Procedure ASSIGN given in Definition 2). This definition depends on whether *v* is the root *r* of *T* or not, as we cannot add an edge to *T* that begins above the root of *T* . Therefore, when *v* ≠ *r*, we let *Pairs*(*v*) denote the set of all ordered pairs of nodes (*u, w*) in *T* such that *lca*_*T*_ (*u, w*) = *v*; note that this allows *u* = *v* with *w* below *v*, and so includes pairs of the form (*v, w*) with *w* below *v*. These pairs (*v, w*) with *w* below *v* imply a ghost lineage [Tricou et al., 2025], which is the result of incomplete taxon sampling. When *v* = *r*, we define *Pairs*(*v*) to be the set of all ordered pairs (*u, w*) in *T* where *lca*_*T*_ (*u, w*) = *r* but neither *u* nor *w* is *r*. Then,

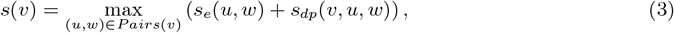

We now show how we calculate *s*_*dp*_(*v, u, w*). Recall that *lca*_*T*_ (*u, w*) = *v* and the assumption is that we are adding the edge *e* above (*u, w*) and hence *v* is the target node. Consider the cycle *γ*_(*u,w*)_ formed by adding the edge *e*. Since a level-1 phylogenetic network *N* cannot have a pair of cycles that share vertices, this means that the target node *x* for any other edge that is added cannot be in this cycle (see Definition 2); otherwise *x* would be in two cycles, contradicting that *N* is level-1. We use this to derive the formula for *s*_*dp*_(*v, u, w*), but note that it depends on whether *v* = *u* (as this changes the cycle). If *v* ≠ *u*, we obtain:

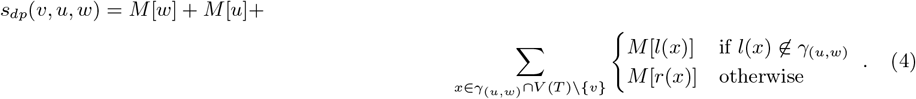

When *v* = *u*, we obtain:

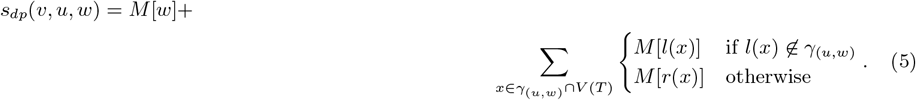

Equations 4 and 5 sum the subproblem scores for all the subnetworks hanging off of γ_(*u,w*)_.

### 2.3 Proof of correctness

We now prove that CAMUS solves its optimization problem exactly.

#### Theorem 1.

*The algorithm gives an assignment of non-tree edges to add the tree T that maximizes the number of quartets in* Q *that are satisfied, under the constraint that we create a level-1 phylogenetic network*.

*Proof*. By design, the solution that CAMUS outputs is a level-1 phylogenetic network *N* that extends *T* . Specifically, consider the target nodes for a pair of added edges defined by the backtracking through the DP matrix. If their target nodes are not in relation (neither above the other), then the cycles they create are vertex-disjoint. If some target node *x* is above another target node *y*, then this case would be handled by Equations 4 and 5, which ensure that the created cycles do not share vertices. It is also easy to see that *M* [*r*] is the number of quartet trees in 𝒬 that are satisfied by *N* and not by *T*, where *r* is the root of *T* .

Hence, it remains to be shown that the score of the network found by CAMUS is the largest possible score of any level-1 phylogenetic network that extends *T* . We will prove that for each node *x* in *T, M* [*x*] is the best score that can be found for any level-1 phylogenetic network all of whose added edges have target nodes in *T*_*x*_. We prove this by strong induction on the size of the leafset below *x*. Our base case is where the size of the leafset is 1, which is when *x* is a leaf; for this case, the score is 0, and is trivially correct. The inductive hypothesis is that *M* [*x*] is correctly computed for all nodes *x* that have fewer than *k* leaves in their subtree, and we now consider a node *y* with *k* leaves in its subtree.

Let *N* be an optimal extension of *T*_*y*_ to a level-1 phylogenetic network. If this optimal extension does not include any added edge with target node *y*, then by the inductive hypothesis, its total score is easily seen to be *M* [*l*(*y*)] + *M* [*r*(*y*)], where *l*(*y*) and *r*(*y*) are the left and right children of *y*, respectively.

If the optimal extension includes an added edge *e* with target node *y*, then *e* is added above the pair (*a, b*), where *lca*_*T*_ (*a, b*) = *y*. We will show that in this case, the score of the optimal extension is given by *s*(*y*), thus establishing correctness.

By Lemma 1, each *s*_*e*_(*a, b*) does not depend on another calculations, and hence is correctly computed during the preprocessing. Additionally, due to Lemma 2, we see that it is sufficient to simply search for one edge that maximizes *s*(*y*); we do not need to consider the possibility of multiple edges with a target of *y*. Also note that for the *s*_*dp*_(*y, a, b*) term, *T*_*y*_ + *e* has a cycle, and since *N* is level-1, all the other cycles in *N* are disjoint from this cycle, which means that the target nodes for all other non-tree edges in *N* are not in the cycle. Therefore, for every other cycle, its target node is below some node in the cycle, and its contribution to the score is handled by *s*_*dp*_, given in Equations 4 or 5. Finally, we note that, due to Lemma 1, it is not possible that any quartets are double-counted when summing up the terms in *s*(*y*) or *s*_*dp*_(*y, a, b*).

Thus, the score of the optimal extension of *T*_*y*_ is identical with how *M* [*y*] is defined, as given in Equation 2, and the inductive hypothesis holds.

### 2.4 Running Time

First, we consider the time to compute *δ*_*T*_ (*q*, (*u, w*)). If we assume that we preprocess the LCA of all pairs of vertices, the leafsets for all vertices, and the depth of every node in the tree, we see that we can calculate *δ*_*T*_ for a single *q* in *O*(1) time (since determining which quartets are induced can be determined by their ordering in the cycle (see discussion in Warnow et al. [2025], Gambette et al. [2012]). We note that the running time for calculating all the LCAs and all the depths is *O*(*n*^2^) and *O*(*n*) respectively, and thus can be ignored in the broader context of the time complexity analysis, as later steps will dominate. Given this, the time to solve *s*(*v*) is *O*(*n*^2^(|𝒬| + *n*)). There are *O*(*n*) subproblems that need to be solved, thus making the total naively *O*(*n*^3^(|𝒬| + *n*)); however, this analysis overestimates the running time as no edge is ever considered in the *s*(·) function more than once over all subproblems. Therefore, the total number of edges maximized over across all subproblems is *O*(*n*^2^), making the runtime *O*(*n*^2^(|𝒬| + *n*)).

### 2.5 Reticulation Restriction

By design, the score (the number of quartets in Q that are satisfied) will never decrease as the number of reticulations increases, and so the objective is to find the “right” number of reticulations. This is a general challenge in phylogenetic network estimation, that we address through an extension of CAMUS. Specifically, we introduce a modification to the dynamic programming algorithm in order to retrieve a set of phylogenetic networks, differing by the number of reticulations. Given the set of phylogenetic networks, one for each number of added edges, it then becomes possible to select a suitable number of edges by examining where the score stops improving substantially. We demonstrate this in Figure 2, which shows that on the 16-taxon datasets, in general selecting the network with just one reticulation (i.e., one extra edge) seems best.

**Figure 2:**
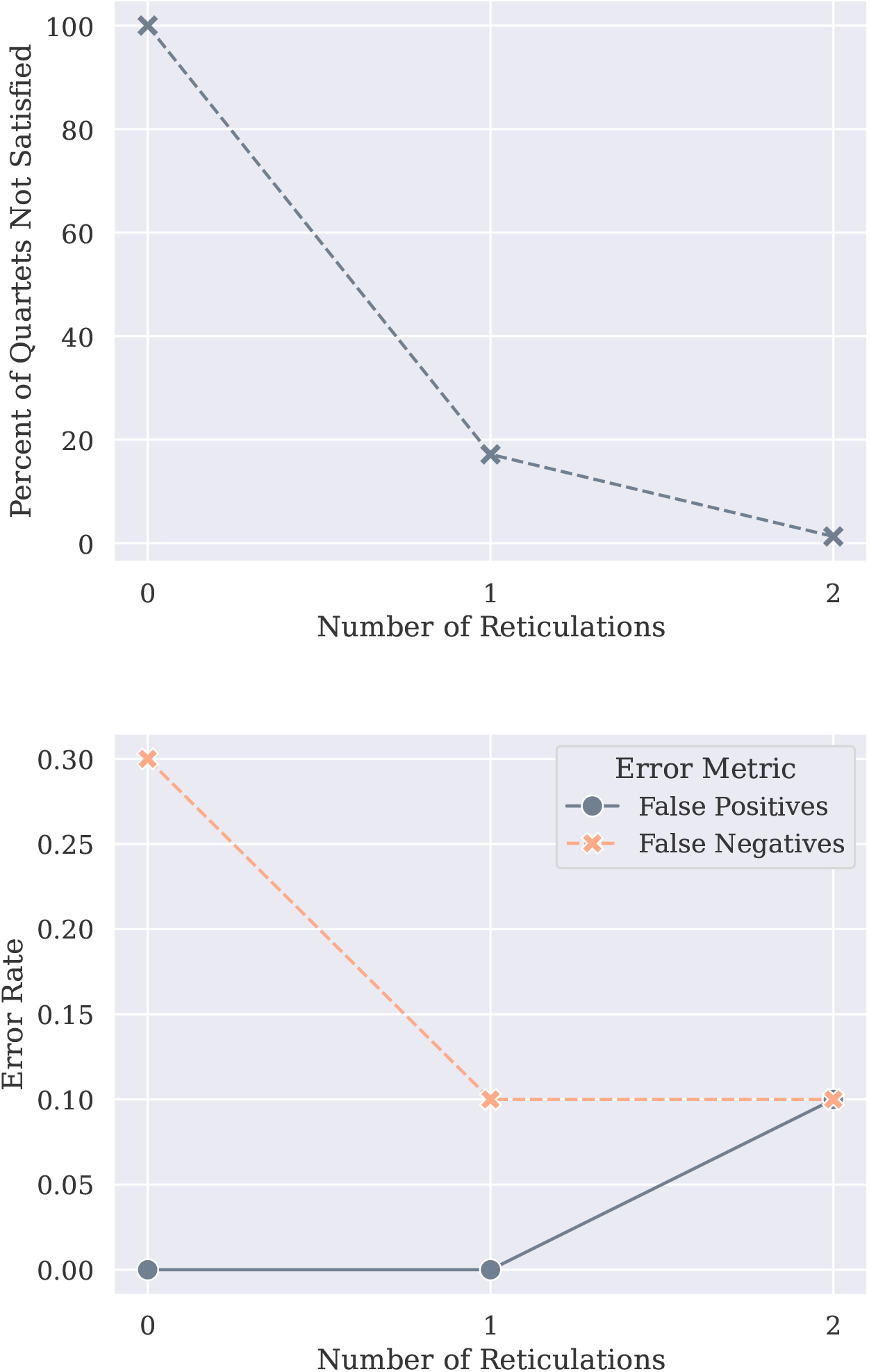
Impact of the number of computed reticulations in the CAMUS network and two error measures on 16-taxon datasets. CAMUS is run using the reticulation restriction mode so that for each number of reticulations, it returns a phylogenetic network optimizing the number of satisfied quartets in 𝒬 with that number of reticulations. Top: percentage of unsatisfied quartet trees from the input multi-set 𝒬. Bottom: phylogenetic reconstruction error as a function of number of reticulations. Based on the improvement in optimization score, one reticulation would be selected, which also corresponds to the most topologically accurate solution for this input.

To this end, we introduce a modification to the algorithm with the following properties. Given a constraint tree *T* and a set of quartets Q, the modification returns *m* networks *N*_1_, *N*_2_, · · ·, *N*_*m*_ so that:

- All networks are level-1.
- All networks contain the constraint tree *T* .
- *N*_*k*_ for 1 ≤ *k* ≤ *m* contains exactly *k* non-tree edges.
- All networks maximize the number of quartets from 𝒬 they contain, subject to the above constraints.

This can be accomplished by a relatively simple modification of the dynamic programming algorithm. Instead of having the dynamic programming lookup table only have *n* entries (where *n* is the number of taxa), we make it a matrix that has *n* rows and *m* columns, where *m* is the number of reticulations after which the score does not improve. However, it is known that a level-1 phylogenetic network on *n* leaves will not have more than *n* − 1 reticulations, so *m* ≤ *n* − 1 is guaranteed. The subproblem *M* [*v, k*] represents the solution found for subtree *T*_*v*_ of the network limited to allowing *k* non-tree edges, and we solve each subproblem in order of increasing *k*, dynamically adding new columns to our matrix, until we no longer get an improvement in score.

Thus, we need to solve each of the *n* original subproblems *O*(*m*) times. It takes *O*(*m*^2^) time to find the correct DP look-ups to solve each problem optimally, thus the running time of the overall algorithm goes from *O*(*n*^2^(|𝒬| + *n*)) to *O*(*n*^2^(|𝒬| + *m*^3^*n*)). However, in our experiments *m << n* and the running time is dominated by the *O*(*n*^2^|𝒬|) term so there is very little running time overhead in practice.

## 3 Pipelines using CAMUS

CAMUS can be used with any constraint tree *T* and set 𝒬 of quartet trees, and the way that each is estimated from the data depends on assumptions about the process and the type of data. We present two pipelines here: one for use with gene trees that evolve down phylogenetic networks under the NMSC and the other for use with SNPs that evolve down phylogenetic networks under a homoplasy-free model described in Gusfield [2005], Warnow et al. [2025].

### 3.1 Pipelines for use with SNPs under Gusfield’s model

Gusfield’s model addresses a population-genetics context where the sites have two states and evolve under the infinite sites assumption, which means that they change state only once during their evolution within the phylogenetic network. Gusfield referred to such sites as SNPs, and proposed a method to construct the rooted level-1 phylogenetic network from such sites, provided that the network does not have small cycles [Gusfield, 2005]. His approach was proven correct when the set of sites covers all the edges in the network and each site changes exactly once in Warnow et al. [2025], which also established that his method, as well as some quartet-based methods, are guaranteed accurate under the same conditions. Moreover, Warnow et al. [2025] proposed a hierarchical model that has gene trees evolving down the network and sites evolving down each gene tree, and proved that Gusfield’s algorithm as well as some quartet-based methods would be statistically consistent under the model. The main limitation with these methods is that when the data are either insufficient or have errors (i.e., evolve with homoplasy), then the methods will just fail to return a network. Thus, these methods produce perfect reconstructions of the network given enough error-free data and otherwise generally fail to return anything,

We now show we can use CAMUS to address this limitation, provided we have an outgroup (which is typically the case). First, we compute quartet trees *ab*|*cd* where *a* and *b* exhibit one state and *c* and *d* exhibit the other state, and we let 𝒬 be the multi-set of all quartet trees found by this technique. According to the theory given in Warnow et al. [2025], if each site changes at most once in the network, then at most two quartet trees will be obtained for any set of four species, with one of the quartet trees matching the underlying tree. Furthermore, as the number of gene trees increases, this set will return *𝒬* (*N*_*r*_) with probability converging to 1. We run ASTRAL on this set of quartet trees to obtain the constraint tree *T* . We then use an outgroup species to root *T*, and run CAMUS on the input pair (rooted version of *T* and multi-set 𝒬).

This pipeline has several nice features. First, after computing the ASTRAL tree, it runs in polynomial time and always returns a phylogenetic network that contains the ASTRAL tree. Second, if the network is tree-based and the most frequently observed quartet tree is induced by the underlying tree, then this pipeline is a statistically consistent estimate of the true network. This can be seen by noting that under these conditions, (1) the ASTRAL tree is a statistically consistent estimate of the underlying tree, and (2) as the number of sites goes to infinity, the set 𝒬 will converge to 𝒬 (*N*_*r*_) (i.e., the set of quartet trees induced by rooted trees contained in the rooted network *N*), as proven in Warnow et al. [2025].

### 3.2 Pipeline for use with gene trees under the NMSC

The input is a set of estimated gene trees and the assumption is that the true gene trees evolve down a level-1 phylogenetic network under the NMSC model, which allows for both ILS and HGT. A simple approach for estimating *T* and 𝒬 is to let *T* be any tree that is computed on the input set of gene trees that we hope is one of the trees contained in the network, and 𝒬 could be all quartet trees induced by any gene tree.

Unfortunately, there is no current method for estimating an underlying tree within the network that is provably statistically consistent [Ané et al., 2024]. However, we can consider methods, such as ASTRAL [Mirarab et al., 2014], to estimate *T* . ASTRAL is based on examination of the quartet tree distribution, and uses the fact that under the MSC the most probable quartet tree has strictly higher probability than the remaining two, which have equal probability. Thus, each gene tree induces 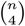 quartet trees (one tree for every set of four leaves), and together the set of gene trees thus defines a distribution on quartet tree topologies that we can use to estimate *T* ^*^ and produce a set 𝒬.

Although ASTRAL can be provably inaccurate at estimating an underlying tree within the network under some conditions [Solís-Lemus et al., 2016, Ané et al., 2024, Dinh and Baños, 2025], some simulation studies have shown it can be highly accurate and also scalable for estimating the underlying tree under a random HGT model [Davidson et al., 2015]. This observation is supported by theory that shows that when HGT is random and not too frequent, that quartet-based methods, such as ASTRAL, are statistically consistent at estimating the underlying species tree [Roch and Snir, 2013].

Given an estimate of an underlying tree *T*, the next step is to estimate 𝒬 from the input data. If we assume that the network is tree-based, with underlying tree *T* ^*^, then gene trees can differ from *T* ^*^ due to both ILS and HGT. In this case, the quartet tree distribution can then be examined to see what additional insight they provide into whether reticulate edges are also needed. Our approach is to take the distribution on quartet trees, and decide whether for a given set of four species, the distribution is suggestive of an ILS-only scenario. If so, then we would just keep the most frequently observed quartet tree, and otherwise we would keep the top two most frequent quartet trees. The motivation for this approach is that we do not want to include quartet trees that are not in *𝒬* (*N*_*r*_), where *N* is the true phylogenetic network. On the other hand, we do not want to eliminate these quartet trees.

In an extreme case, there may be little to no HGT, which would be indicated by finding that the distribution on quartet trees defined by the gene trees fits the multi-species coalescent (MSC) model [Takahata, 1989] under which, for every four species, the most probable quartet tree matches the species tree and the other two quartet trees have the same lower probability. Thus, to the extent that for a set of four species, we get a pattern that matches the MSC, we might decide to not include *any* quartet tree in 𝒬 beyond the most frequently observed quartet tree. We use this idea in designing a simple method for filtering the set 𝒬_0_ of 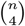 quartet trees to a smaller set that seems reflective of HGT rather than only ILS.

#### Quartet Filtering

Given a threshold *t* ∈ [0, 1], we filter the set Q_0_ of all possible quartet trees as follows. For a given set of four species *a, b, c, d* and quartet topologies *q*_1_, *q*_2_, and *q*_3_, we let *f*_*i*_ denote the relative frequency of *q*_*i*_ (i.e., proportion of genes exhibiting of *q*_*i*_), and we assume without loss of generality that 1 ≥ *f*_1_ ≥ *f*_2_ ≥ *f*_3_ ≥ 0. We always keep the most frequently observed quartet tree *q*_1_ and we do not include *q*_3_. However, we *also* include *q*_2_ if the gap between *f*_2_ and *f*_3_ is large enough, as we define in Quartet Filtering, given in Algorithm 1. In the end, we return the set of quartet trees, denoted by 𝒦.

Now, consider the case where the true evolutionary history is tree-like (i.e., does not require any additional edges) so that the gene trees evolve down the tree under the MSC, which allows for ILS but not HGT. In this case, for large numbers of gene trees, we expect the distribution on quartet tree topologies to be close to the expected values under the MSC, which is *f*_1_ *> f*_2_ = *f*_3_. For finite numbers of gene trees, we would not expect *f*_2_ = *f*_3_, but as the number of genes increases, the gap between *f*_2_ and *f*_3_ will approach 0.

We use a simple technique for building 𝒬: we always include *q*_1_ but we decide whether to include *q*_2_ in the set 𝒬 as follows: if *f*_2_ − *f*_3_ *> t* · (*f*_2_ + *f*_3_), then we include *q*_2_ and otherwise do not. Note that for *t* small enough (specifically,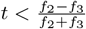), then we will include *q*_2_. We formalize this in the *Quartet Filtering* algorithm in Algorithm 1.

##### Algorithm 1 Quartet Filtering

The input is a multi-set of quartet trees on four species, and the output is a multi-set, including the copies of the most frequently observed quartet tree or the copies of the top two most frequent. This is applied to all sets of four species to determine the multiset 𝒬 given as input to CAMUS.

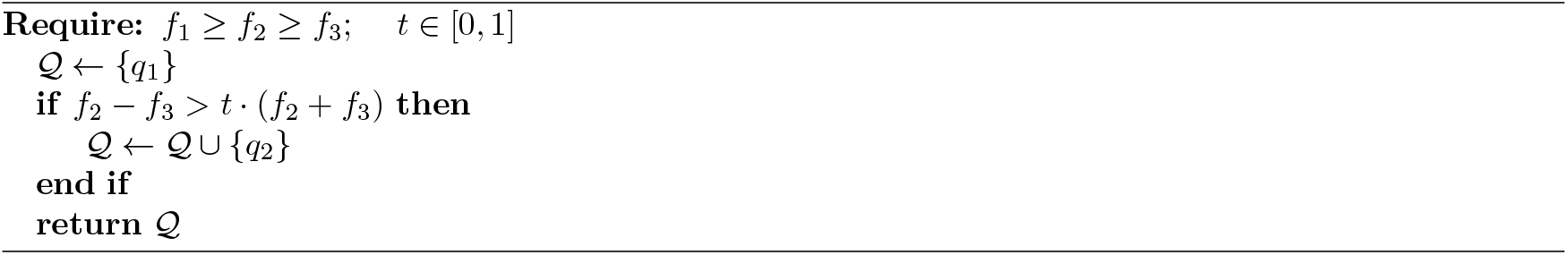

Note the impact of *t* on Quartet Filtering. How we set *t* determines which quartet trees are used to find edges to add to *T* . Even if *T* is an induced tree in the true network *N*, if we let *t* be too small, we may include many quartet trees, which could produce a network that is more complex than the true network; in contrast, if we let *t* be too large, then we may not include enough quartet trees, which would produce a network that is less complex than the true network. Therefore, in Experiment 1, we used algorithm design data (separate from the testing data used in Experiments 2 and 3) to pick a default value for *t*, and then used that value in Experiments 2 and 3.

## 4 Experimental Study

We evaluate CAMUS in comparison to other methods for estimating level-1 phylogenetic networks with respect to topological accuracy and computational performance on synthetic datasets that evolve down level-1 phylogenetic networks under the NMC; this enables an evaluation of accuracy, since the true phylogenetic network on a biological dataset is not known with certainty.

### 4.1 Methods

We compare CAMUS to SNaQ and PhyloNet-MPL [Yu and Nakhleh, 2015], two leading methods for phylogenetic network estimation that also take as input a set of estimated gene trees and return level-1 phylogenetic networks. The choice of PhyloNet-MPL and SNaQ is based on their frequency of use in biological dataset analyses and that they are able to complete on datasets with at least 10 species. We gave the same set of estimated gene trees to each method, as well as the same tree *T*, which was treated either as a constraint tree (for one way of running PhyloNet-MPL) or just as a starting tree (for SNaQ and the other way of running PhyloNet-MPL).

While SNaQ takes as input unrooted gene trees, PhyloNet-MPL requires rooted gene trees; we enable this through the use of the outgroup species in each dataset. While SNaQ always allows the input tree to change, PhyloNet-MPL can be run in two ways: allowing the tree to change or considering it fixed throughout the search. We refer to the second way of running PhyloNet-MPL as “PhyloNet-MPL(FT)”, where “FT” stands for “Fixed Tree”.

### 4.2 Synthetic Datasets

We generated simulated datasets with 1000 gene trees that evolve down level-1 phylogenetic networks. The datasets used in the Algorithm Design (Experiment 1) are disjoint from the datasets used to compare CAMUS to PhyloNet-MPL and SNaQ in Experiments 2 and 3. These datasets range from 16 species to 201 species, and each dataset has an outgroup.

To produce our simulated datasets, we use SiPhyNetwork [Justison et al., 2023] to create a simulated true phylogenetic network. These networks are then filtered to ensure that they are level-1, and an outgroup taxon was added, so that the rooting can be determined in later steps. After that, we used PhyloCoalSimulations [Fogg et al., 2023] to produce true gene trees under the NMSC; thus, these gene trees have discordance from both Incomplete Lineage Sorting (ILS) as well as the HGT events. To produce estimated gene trees, we then evolved DNA sequences of length 500 down the trees with INDELible [Fletcher and Yang, 2009] under GTRGAMMA, and estimated trees with IQ-TREE 3 [Wong et al., 2025] (for the 16–26 taxon datasets) and FastTree 2 [Price et al., 2010] (for the 26–201 taxon datasets). This results in gene trees with an average estimation error of 20% in the case of IQ-TREE and 21% in the case of FastTree 2. We used Ultrafast Bootstrap Approximation with IQTREE [Hoang et al., 2018] in order to estimate branch support. Our method requires a constraint tree for estimation, and PhyloNet-MPL and SNaQ can be assisted by a starting network (or tree), thus we estimate this tree with ASTRAL 4 [Mirarab et al., 2014, Zhang et al., 2025] using the estimated gene trees. More details about the parameters and exact commands used for all of the above are provided in the Supplementary Materials.

### 4.3 Computational Environment

We ran all methods with a time limit of 12 hours and a memory limit of 256 GB on the University of Illinois Urbana-Champaign Campus Cluster.

### 4.4 Evaluating topological error

To assess phylogenetic network topology estimation error, we use the cluster metric from PhyloNet’s CmpNets command [Nakhleh and Wang, 2005], which calculates a false positive and false negative rate based on soft-wired clusters. We were able to use this code on networks with up to 51 species but not larger, due to computational limitations (i.e., more than 5 hours to score a single 51-species network). Therefore we calculate this error rate only for the networks with at most 51 species.

### 4.5 Experiments

Our evaluations are split into three experiments:

- **Experiment 1**. Designing the pipeline for phylogenetic network estimation from gene trees using CAMUS; this involves setting the default value for threshold *t* in Quartet Filtering that we then use in subsequent experiments.
- **Experiment 2**. Evaluating computational performance of CAMUS, PhyloNet-MPL, and SNaQ on datasets with 16 to 201 species.
- **Experiment 3**. Evaluating accuracy of CAMUS, PhyloNet-MPL, and SNaQ on datasets with 16 to 51 species.

For all experiments, we show results on 20 replicates containing 1000 gene trees each. Experiment 1 uses a different set of replicates on 26 taxa than Experiments 2 and 3.

#### Evaluation

We evaluate runtime and topological error for SNaQ, PhyloNet-MPL, PhyloNet-MPL(FT), and CAMUS, each given the ASTRAL tree as either a constraint tree (for PhyloNet-MPL(FT) and CAMUS) or as a starting tree (for SNaQ and PhyloNet-MPL). In addition, each method is used to create a level-1 phylogenetic network with exactly one reticulation; while it is possible that allowing more reticulations would improve accuracy, selecting the correct number of reticulations is non-trivial from a statistical viewpoint and often performed using a visual inspection such as provided in Figure 2. Furthermore, it would require running SNaQ, PhyloNet-MPL, and PhyloNet-MPL multiple times (for each possible number of reticulations), which is computationally infeasible for these methods (unlike for CAMUS that automatically provides this in its default setting, using the Reticulation Restriction modification).

## 5 Results

### 5.1 Experiment 1

Recall that when we estimate the phylogenetic network from gene trees, we construct a multi-set 𝒬 of quartet trees that will contain some but generally not all the quartet trees found in the input gene trees. The selection of which quartets to include is described above in the section labeled “Quartet Filtering”, whose output depends on a parameter *t*. In Experiment 1, we assess the impact of *t* on the final accuracy of the estimated level-1 phylogenetic network. All results are shown on 20 replicates of the 26-taxon dataset (25 taxa plus 1 outgroup taxon) with 1000 estimated gene trees.

As seen in Figure 3, we see that applying a threshold *t >* 0 (i.e., not including all possible quartet trees) improves accuracy, and that *t* = 0.5 provides the best accuracy of the tested thresholds. Henceforth, we use *t* = 0.5 for all further experiments.

**Figure 3:**
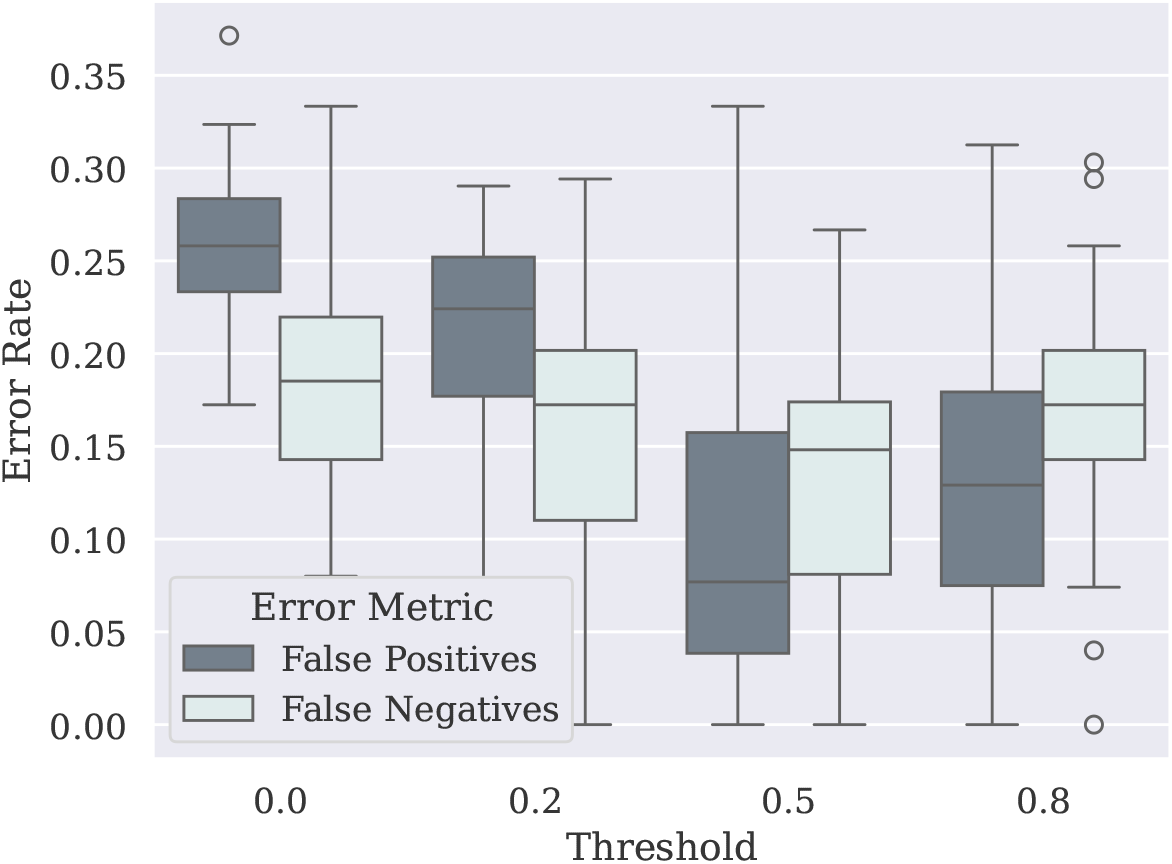
Impact of quartet filtering threshold on phylogenetic network error rate computed by CAMUS. Results are shown on the algorithm design datasets with 26-taxon FastTree gene trees where CAMUS adds one edge to produce 1 reticulation; we report the cluster metric error rate. Setting the threshold *t* = 0.5 produces the lowest error.

### 5.2 Experiment 2

Experiment 2 explores the differences between the computational performance of our selected methods. We examine several different numbers of taxa, between 16 to 201, each with 1000 estimated gene trees, across 20 replicates. In this experiment, CAMUS was run using the Reticulation Restriction mode, where it computes the best setting for all numbers of added non-tree edges (up to when the score stops improving), while the other methods were run only long enough to find the best network with one non-tree edge.

As seen in Figure 4 (left)), all methods except CAMUS fail at some point to complete within our time limit of 12 hours, before reaching the largest dataset on 201 taxa. SNaQ is only able to run on our smallest number of species, failing on 26 taxa; PhyloNet-MPL is a bit faster, but times out on 51 taxa; and PhyloNet-MPL(FT) manages to run on 51 taxa, but cannot complete on the 101-taxon dataset. Moreover, the running time for CAMUS on 16-species datasets was under a second, and increased with the number of species but was still under a minute when analyzing 51-species datasets. CAMUS runs in roughly 20 minutes for 101 species, and completes on datasets with 201 species in just 2–3 hours.

**Figure 4:**
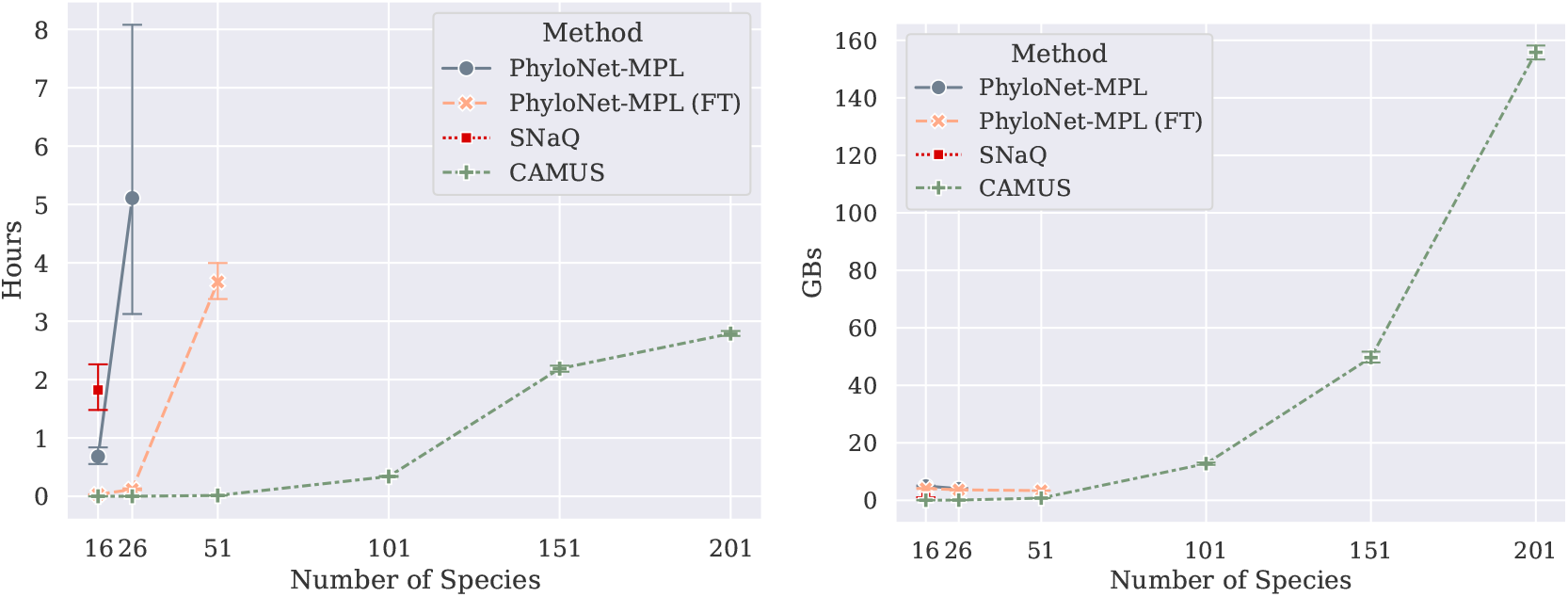
Computational performance of methods as a function of the number of species given 1000 estimated gene trees. Left: running time; right: peak memory usage. Methods do not appear if they do not complete within 12 hours.

We also looked at the memory usage for the methods (Figure 4 (right)). Memory usage was relatively low for all methods on the datasets with at most 51 species, and CAMUS used less memory than the other methods on these datasets. However, the memory usage of CAMUS increased with the number of species, reaching 156GB for 201 species (a condition the other methods could not analyze within the allowed time). This increase in memory usage is unsurprising, as CAMUS memory usage scales linearly with the number of quartets (i.e., *O*(*n*^4^)).

### 5.3 Experiment 3

This experiment evaluates all methods on the testing datasets for topological accuracy.

All methods complete within 12 hours on 16 species (Figure 5 (left)), and overall SNaQ had the worst accuracy, while PhyloNet-MPL without a fixed tree was the most accurate. CAMUS was slightly less accurate than PhyloNet-MPL without a fixed tree, and was more accurate than PhyloNet-MPL(FT).

**Figure 5:**
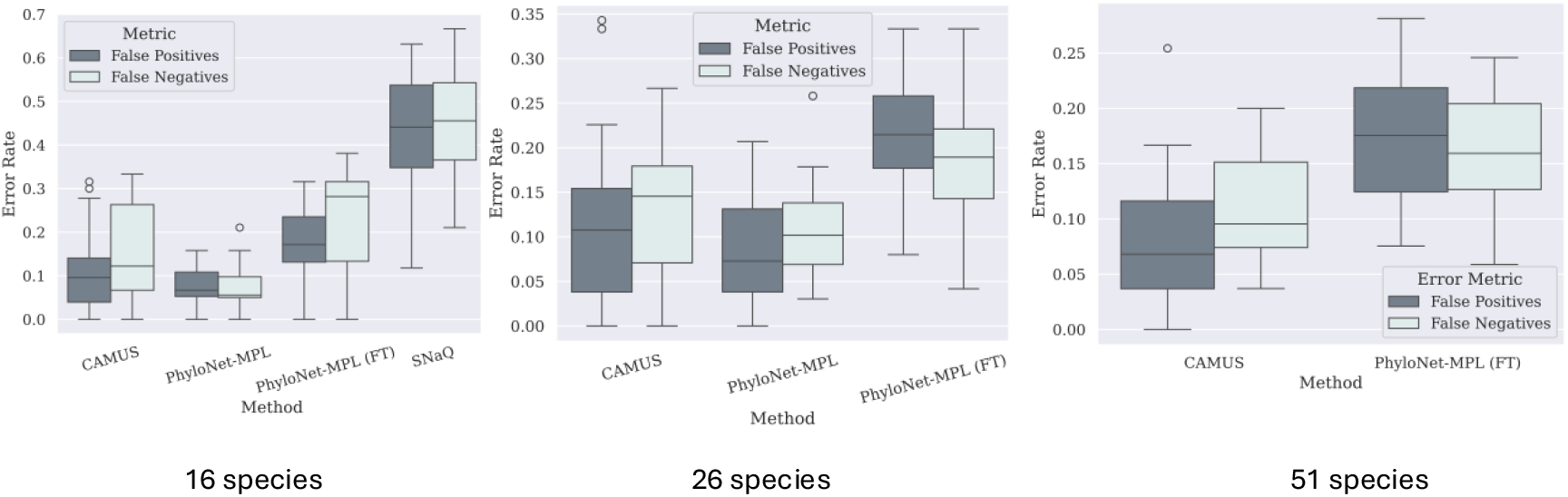
Phylogenetic network estimation error. We show error rates (false negative and false positive) for SNaQ, PhyloNet-MPL, PhyloNet-MPL(FT), and CAMUS on simulated datasets. Left: 16-species datasets; middle: 26-species datasets; right: 51-species datasets. Each condition has 1000 estimated gene trees and 20 replicates. Results not shown for a particular number of species indicate the method failed to complete within the allotted time (12 hours) given 256 GB of memory. Each phylogenetic network estimation method creates a level-1 phylogenetic network with one reticulation, and uses the ASTRAL tree either as a constraint tree or as a starting tree (this is true for all experiments in this study).

On 26 species (Figure 5 (middle)), SNaQ could not complete within the allowed time, but the remaining methods did complete. On these datasets, again PhyloNet-MPL without a fixed tree had the best accuracy, followed by CAMUS, and then by PhyloNet-MPL(FT).

Finally, on 51 species (Figure 5) (right)), only CAMUS and PhyloNet-MPL(FT) completed within the allowed time and CAMUS had much better accuracy than PhyloNet-MPL(FT).

## 6 Conclusion

We have presented CAMUS, a polynomial time method that takes as input a rooted binary tree *T* and a set 𝒬 of quartet trees, and returns a level-1 phylogenetic network that satisfies as many of these quartet trees in 𝒬 as possible. CAMUS can be used within pipelines to estimate phylogenetic networks, and we described two such pipelines (one for Gusfield’s model, which is relevant to population genetics, and the other for estimating phylogenetic networks from gene trees under the NMSC). These pipelines differ, and demonstrate the flexibility of CAMUS, where the tree *T* and set 𝒬 can be defined as suited for the input dataset and model of evolution. Thus, CAMUS is a flexible tool for use within pipelines for phylogenetic network estimation, which will allow it to be used in new pipelines with ease.

Our simulation study explored CAMUS specifically for estimating level-1 phylogenetic networks under the NMSC, where the input is a set of estimated gene trees. We showed that CAMUS has much greater speed and scalability than PhyloNet-MPL (used with or without a fixed tree) and SNaQ. CAMUS was superior in accuracy to SNaQ and PhyloNet-MPL used with a fixed tree, but was slightly less accurate than PhyloNet-MPL used without a fixed tree. Furthermore, CAMUS successfully completes on datasets with 201 species and 1000 gene trees, while none of the other methods was able to complete within 12 hours on the dataset with 101 species. Furthermore, the most accurate method, PhyloNet-MPL, fails to complete on the inputs we studied with more than 26 species. Thus, CAMUS is a practical tool for accurate phylogenetic network estimation, scaling to at least 201 species.

This study suggests several directions for future research, but due to space limitations, we mention only a few. Given the improvement in accuracy for PhyloNet-MPL when it allows the input tree to change, using CAMUS within a heuristic search to allow it to explore different constraint trees could improve accuracy and maintain scalability. Furthermore, we have started to explore the impact on accuracy of collapsing low support edges in the gene trees, and our initial investigations on a limited sample suggest that this improves accuracy. We will also explore ways to reduce the runtime, which would include not computing all the quartet trees; such an approach could be based on “short quartets”, as studied in several papers, starting with Erdős et al. [1999]; this could greatly improve scalability since typically there are *o*(*n*^4^) short quartets. Finally, we plan to explore CAMUS on biological datasets, but because there are no biological datasets where the ground truth phylogenetic network is known, evaluating these results will require additional expertise.

## 7 Data availability

CAMUS, implemented in Go, is available in open-source form on GitHub [Willson, 2025] and all the data generated for this study are freely available at Willson and Warnow [2026]. CAMUS utilizes Gotree [Lemoine and Gascuel, 2021] as a core dependency.

