## Supplementary Materials for "CAMUS: Scalable Phylogenetic Network Estimation"

### 1 Data Simulation

The following pipeline is used for data simulation

- Generate model networks using SiPhyNetworks [1].
- Outgroup taxa are then added to these networks.
- Gene trees are then generated from these networks using PhyloCoalSimulations [2].
- Sequences are generated from these gene trees using INDELible [3].
- FastTree2 [4] or IQTree3 [5] is then used to generate estimated gene trees.

#### 1.1 Model Networks

Model networks are generated from $n$ taxa, where $n \in \{15, 25, 50, 100, 150, 200\}$ (plus one outgroup). SiPhyNetwork (v1.1.0) has the following additional parameters: speciation rate $\lambda$, hybridization rate $\mu$, and extinction rate $\nu$, as well as a vector `hyperprobs` controlling the probability that lineages are generative, degenerative, or neutral. A function to determine inheritance proportions is also taken as input. 50 replicates are generated for all model conditions.

$\lambda = 1$ for all model conditions; `hyperprobs` $= \langle 0.5, 0.25, 0.25 \rangle$ where the values are lineage generative, lineage degenerative, and lineage neutral, respectively. Inheritance proportions are drawn from a Beta(10,10) distribution.

In addition, the following filtering criteria were in forced: each network must be level-1, must have at least one reticulation, and must have exactly $n$ taxa.

Both $\nu$ and $\mu$ were adjusted (primarily to make finding a set of model networks computationally feasible as both of these parameters affect the likelihood of a network being level-1).

- $n = 25$, $\nu = 0.2$, $\mu = 0.025$
- $n = 50$, $\nu = 0.2$, $\mu = 0.0025$

- $n = 100$, $\nu = 0.05$, $\mu = 0.0025$
- $n = 150$, $\nu = 0.005$, $\mu = 0.0025$
- $n = 200$, $\nu = 0.005$, $\mu = 0.001$

Next, an outgroup taxa was added. A single vertex was created as the root, then the outgroup leaf was added with an edge length $[0.9, 1.0]$ (uniform distribution); finally, the previously generated network was attached to the other side of the root on the other side of a branch with length $[0.0, 0.1]$.

To generate the networks, we used an R script, `generate-networks.R` and then add outgroups to these networks with a python script `add-outgroup.py`, using the following commands.

```
Rscript generate-networks.R $ntaxa 50 $nu $mu
find "n$ntaxa" -name "true_net.nwk" -exec python add-outgroup.py {} \;
```

### 1.2 Gene Trees

True gene trees were simulated from the model networks using PhyloCoalSimulations (v1.0.0) using default settings (i.e., single individual per species). Then sequences are generated with INDELible. All sequences are generated a with length of 500 bp. This resulted in trees with $\approx 21\%$ gene tree estimation error.

To run INDELible we used the same process and scripts as in [6] (see that paper's Supplementary Materials for more details). We have included a table with the parameters used below (Table 1).

| Parameters | Values |
|---|---|
| Base Frequencies | Dirichlet(T=113.48869,C=69.02545,A=78.66144,G=9983793) |
| Transition Rate | Dirichlet(CT=12.776722,AT=20.869581,GT=5.647810, AC=9.863668,GC=30.679899,AG=3.199725) |
| Gamma rate variation | Log-normal($\mu$-0.470703916, $\sigma$=0.348667224) |

Table 1: INDELible Parameters

We used FastTree (v2.1.11) to estimate gene trees from the sequences generated using INDELible, using the command:

```
fasttree -nt -gtr $input > $output
```

In the cases where IQTree3 is used, we use the command:

```
iqtree3 -T 32 -s $input -m GTR+G -bb 1000
```

#### 1.3 Network Estimation

The following command was used to run ASTRAL-IV [7] (v1.23.4.6) for the starting/constraint networks.

```
astral4 -t 32 -o $output $input 2> $log
```

Trees are then rerooted on the outgroup using the python script `root-outgroup.py` as so.

```
python root-outgroup.py $tree
```

`root-outgroup.py` uses TreeSwift [8] as a dependancy.

##### 1.3.1 CAMUS

CAMUS was run with the following command

```
camus -n 32 -q 2 -t $threshold -o $output $const_tree $gene_trees
```

Older versions of CAMUS required `-q 2` to enable quartet filtering. This is now turned on by default and thus is unnecessary.

##### 1.3.2 SNaQ

For SNaQ (v1.1) [9], we used `countquartetsintrees(·)` from PhyloNetworks [10] to calculate the quartet concordance factors, then we pass these concordance factors directly to SNaQ. We use the script `run_snaq.jl` to achieve this. We call this script with the command

```
julia --threads=auto run_snaq.jl $gene_trees  \
    $start_tree $output 1 32
```

##### 1.3.3 PhyloNet-MPL

For PhyloNet-MPL (v3.8.2) [11, 12] we used a Python wrapper script to create the nexus files: `run_phylonet.py`. Then, to run PhyloNet-MPL we used the command

```
python run_phylonet.py \
    -p PhyloNetv3_8_2.jar \
    -g $gene_trees \
    -s $start_tree \
```

```
    -r 1 \
    -n 32 > $log 2>&1
```

And for PhyloNet-MPL (FT) we add the `-f` argument to the script.
